# Self-Assembling Synthetic Peptides Remodel Mitochondrial Properties via Nanopore Formation

**DOI:** 10.1101/2025.08.26.672490

**Authors:** Nhung Thi Hong Nguyen, Shoko Fujita, Nanami Takeuchi, Ryuji Kawano, Takanari Inoue

**Affiliations:** Department of Cell Biology, Johns Hopkins University School of Medicine, Baltimore, MD 21205, USA; Center for Cell Dynamics, Johns Hopkins University School of Medicine, Baltimore, MD 21205, USA; Department of Biotechnology and Life Science, Tokyo University of Agriculture and Technology (TUAT) 2-24-16 Naka-cho Koganei-shi Tokyo 184-8588 Japan

## Abstract

Pore-forming peptides (PFPs) are powerful tools for engineering biological membranes *in vitro* and in living cells. They can be derived from natural sources or designed *de novo*, with synthetic peptides offering distinct advantages in rational design unconstrained by natural sequences. While native PFPs have been targeted to specific organelles to modulate their functions, the potential of synthetic PFPs for organelle engineering remains largely unexplored. Here, we show that the 28-amino acid synthetic peptides, SVG28 and its variant SVG28D2, can be selectively targeted to mitochondria for functional alteration. More specifically, molecular dynamics simulations first confirmed the structural stability of the SVG28 nanopore even when fused to the TOM20 transmembrane domain, a mitochondrial targeting sequence. In isolated lipid bilayers, TOM20-SVG28 formed pores with a 0.3-nS, conductance, or 1.0-nm pore. The expression efficiency was further enhanced by two amino acid substitutions (SVG28D2). In cells, both TOM20-SVG28 and TOM20-SVG28D2 localized to the mitochondria, induced morphological alterations, and reduced both membrane potential and ATP production. These observations *in silico, in vitro* and *in cellulo* collectively indicate that the synthetic SVG28 peptides self-assemble into pores in the outer mitochondrial membrane, likely permitting proton permeation. Importantly, the matrix protein Su9 remained confined within mitochondria, suggesting preserved inner membrane integrity, and thus specific effect on the target. This work establishes a framework for the design and application of genetically-encoded synthetic PFPs in living cells, opening new avenues for cellular engineering.

## Introduction

Mitochondria are essential organelles responsible for ATP production, metabolism regulation, and apoptosis induction ^1-5^. The functionality of mitochondria relies on the integrity and dynamic remodeling of their membranes. These membranes play a critical role in preserving the electrochemical differences necessary for energy production and in organizing distinct biochemical activities within the mitochondrion ^6-8^. Disruptions in mitochondrial membrane structure or morphology can affect how mitochondria work, which may cause or exacerbate health problems such as neurodegenerative diseases, cancer, and metabolic disorders ^9-11^.

Pore-forming peptides (PFPs) represent a class of molecules that can integrate into lipid bilayers to trigger formation of pores, which can regulate the selective permeability of ions, signaling molecules, and macromolecules ^12-14^. Natural and synthetic PFPs have been extensively investigated for their capacity to alter membrane properties under *in vitro* conditions ^15-18^. For example, synthetic transmembrane peptides based on the pore-forming domain of TRPV4 can oligomerize to form potassium-selective channels that modulate membrane potential in vascular smooth muscle cells^19-22^. Poly-arginine (Arg_9_) peptides are known for their cell-penetrating abilities and can induce formation of transient pores that in turn facilitate molecular transport ^23; 24^. Voltage-dependent anion channels (VDACs), referred to as mitochondrial porins, are endogenous mitochondrial pore-forming proteins. VDACs modulate metabolite flux across the outer mitochondrial membrane and influence apoptotic pathways ^25-28^.

Despite their structural and functional diversity, the potential of synthetic PFPs engineered to selectively target mitochondria and modulate organelle integrity remains largely unexplored. This critical gap in knowledge is whether these molecules can be directed to mitochondria in living cells to precisely control membrane permeability without triggering catastrophic rupture. Overcoming this challenge through advances in design, targeting strategies, and functional regulation will be transformative for organelle engineering, disease modeling, and therapeutic innovation. In this study, we investigated if and how two artificially designed peptides, SVG28 and SVG28D2, previously demonstrated to form nanopores in planar lipid membranes without membrane rupture ^29; 30^, can be targeted to an intracellular organelle for altering its properties.

## Results

### Designing pore-forming peptides targeted to mitochondria

To demonstrate synthetic PFPs can function in a living-cell context, we aimed to target SVG28 synthetic peptides to mitochondria. Toward this end, the transmembrane domain of TOM20 that is known to target proteins to the mitochondrial outer membrane was fused to the N-terminus of SVG28 (**Fig. 1a**). A Flag tag was inserted with a flexible linker between TOM20 and SVG28 to enable antibody-based detection in cellular experiments without compromising functionality of TOM20 or SVG28.

**Figure 1.**
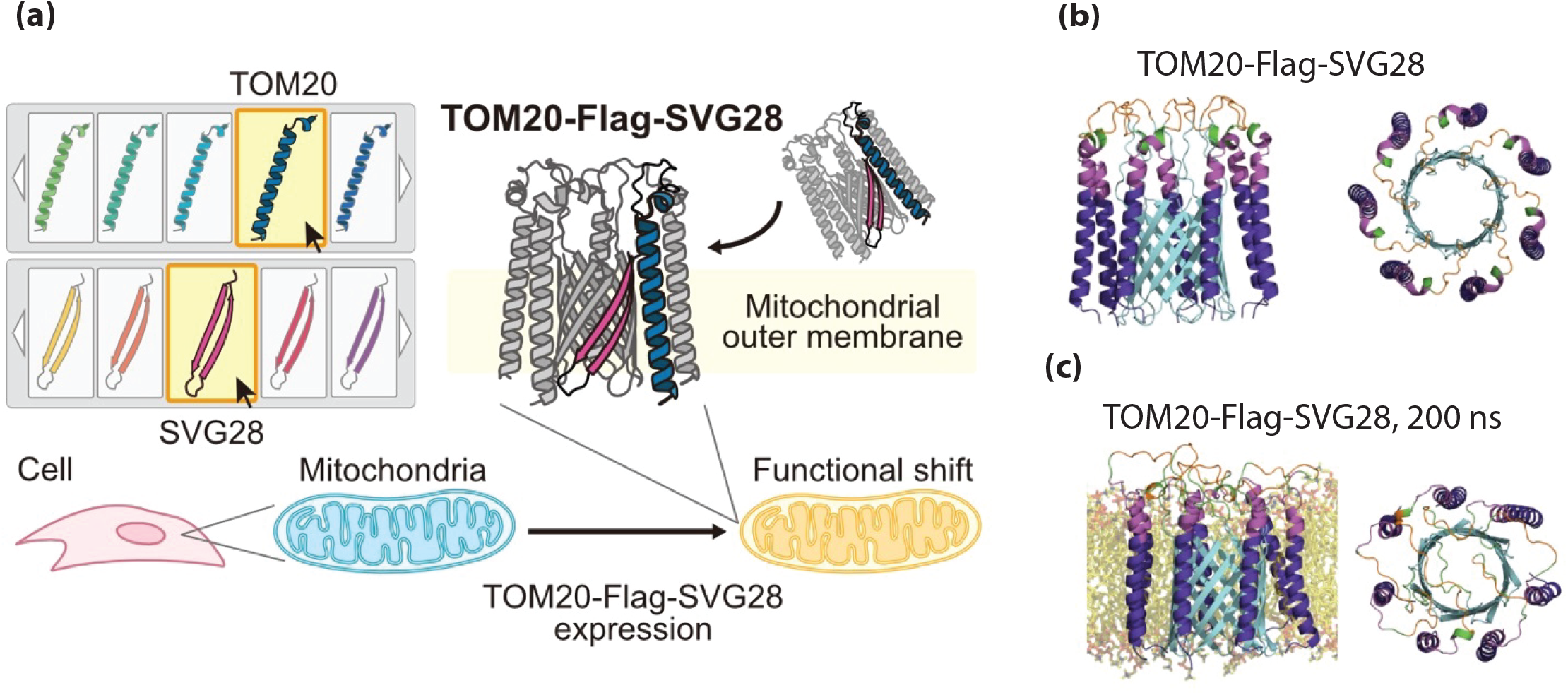
TOM20-Flag-SVG28 conjugate. **(a)** Schematic of the conjugation of TOM20 and SVG28 for functional modulation of mitochondria **(b)** TOM20-Flag-SVG28 structures predicted by *in silico* methods. TOM20 in the transmembrane region is shown in purple; TOM20 outside the membrane in magenta; flag tag in orange; the linkers in green; SVG28 in cyan **(c)** TOM20-Flag-SVG28 pore structures after 200 ns simulation in DOPC:Chol membrane. DOPC and cholesterol molecules are shown in yellow

We first predicted the structures of TOM20-Flag-SVG28 using AlphaFold3 ^31^ and evaluated its stability through all-atom molecular dynamics (MD) simulations. TOM20-Flag-SVG28 showed a configuration similar to native TOM20 ^32^, which has a transmembrane region ranging from residue 1 to 24 (**Fig. 1b**). The TOM20-Flag-SVG28 pore structure was maintained after a 200 ns simulation within 1,2-dioleoyl-sn-glycero-3-phosphocholine:cholesterol (DOPC:Chol, 2:1 molar ratio) bilayer lipid membrane (**Fig. 1c**). While the SVG28 region maintained stable pore structures, the transmembrane TOM20 α-helix modestly fluctuated (**Supplementary Fig. 1a**,**b**). Thus, addition of the TOM20 targeting motif does not appear to impact the overall structural integrity of the SVG28-formed pore. Lastly, the Flag tag was exposed to the water layer as expected (**Fig. 1c**).

### Stability of pore-forming peptides synthesized *in vitro*

To assess the pore-forming capability of TOM-Flag-SVG28 on membrane, we performed electrophysiological measurement across a planar lipid bilayer composed of DOPC:Chol. The molecular mass of the synthesized construct was analyzed by MALDI-TOF mass spectrometry and found to match the expected value (**Supplementary Fig. 1c**), confirming the molecular identity. Upon electrophysiological measurement of TOM20-Flag-SVG28, only five current events were observed during 8 h of recording, indicating that the events occurred infrequently. This low frequency was likely attributed to the poor expression of SVG28, likely due to its highly hydrophobic nature. Therefore, we employed a more hydrophilic variant, SVG28D2 ^29^. TOM20-Flag-SVG28D2 was confirmed to be stable in MD simulation, and was successfully produced using cell-free synthesis followed by MALDI-TOF/MS confirmation (**Supplementary Fig. 2a-c**). TOM20-Flag-SVG28D2 produced approximately a five-fold higher event frequency than TOM20-Flag-SVG28. Based on these findings, subsequent analyses were conducted using both TOM20-Flag-SVG28 and TOM20-Flag-SVG28D2.

### Pore properties of SVG28 peptides in lipid membranes

Initial trials of electrophysiological measurements using lipid bilayers did not indicate stable pore formation, despite the detection of interactions between these molecules and the lipid membranes (**Supplementary Fig. 3**). Suspecting that the presence of transmembrane TOM20 increased unwanted aggregation of SVG28 peptides, we added DOPC:DOPG liposomes to a cell-free protein synthesis solution for efficient membrane insertion of SVG28 peptides with proper folding. As a result, clear pore formation was observed for both TOM20-Flag-SVG28 and TOM20-Flag-SVG28D2 (**Fig. 2a**). To quantitatively evaluate pore-forming ability, the observed current signals were classified into stable or unstable pore formation based on their waveforms. Both constructs exhibited stable pore formation (**Fig. 2b**), with TOM20-Flag-SVG28 displaying a more stable pore formation compared to TOM20-Flag-SVG28D2 (**Fig. 2b**). This is consistent with the observation that SVG28D2, in the absence of TOM20, exhibited less stable pore formation than SVG28. Although direct comparison of the pore-forming stability between these constructs is complicated by factors such as synthesis yield and the extent of liposome incorporation, these results demonstrate that SVG28 peptides are capable of self-assembling into stable pores, even with TOM20 modification.

**Figure 2.**
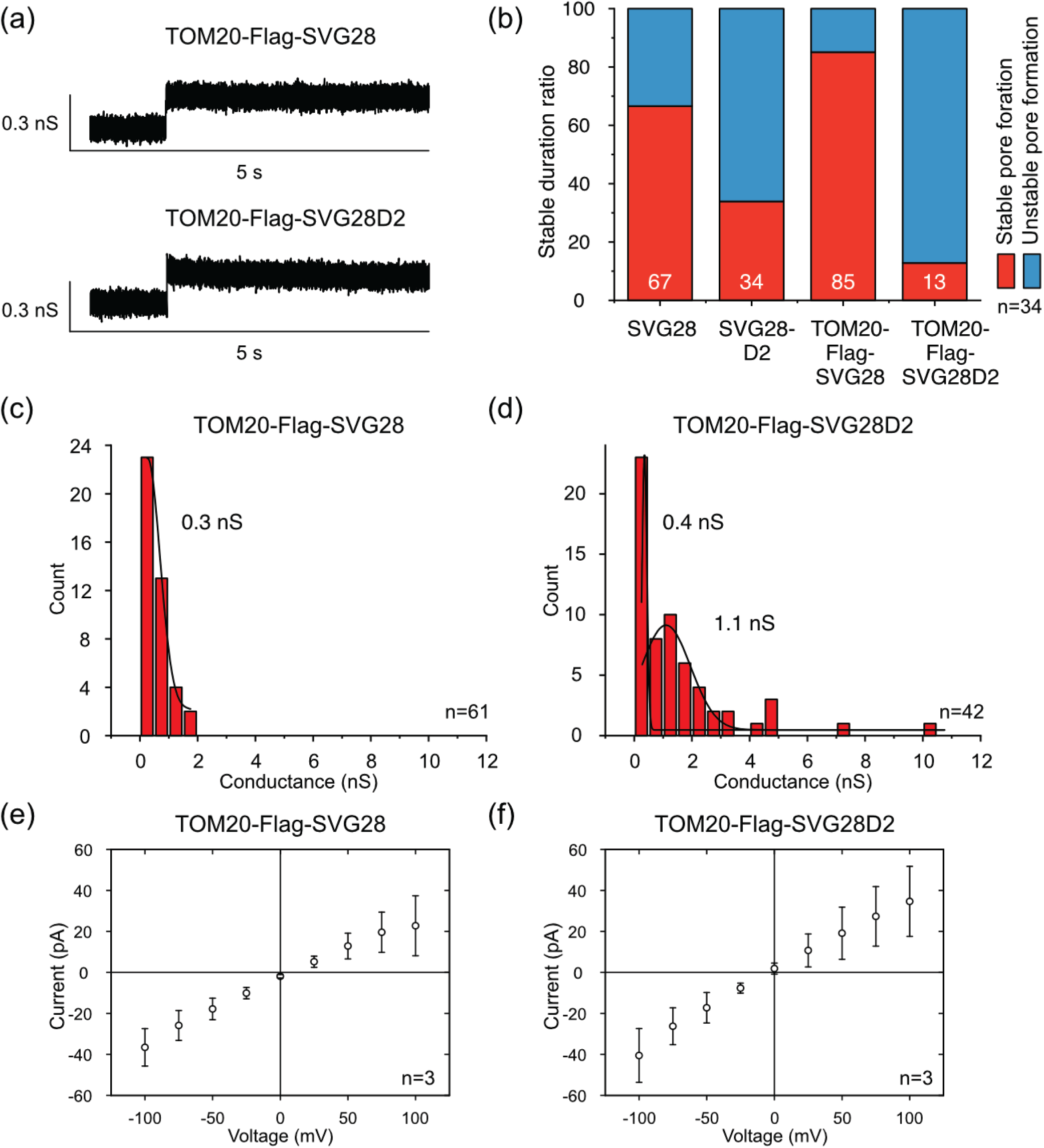
TOM20-Flag-SVG28 and TOM20-Flag-SVG28D2 nanopore characterization. **(a)** The typical current-time traces of TOM20-Flag-SVG28 and TOM20-Flag-SVG28D2 **(b)** The ratio of duration of stable pore formation and unstable pore formation of SVG28, SVG28D2, TOM20-Flag-SVG28, and TOM20-Flag-SVG28D2. For the data of SVG28, the raw data reported in S. Fujita *et al*. (unpublished work) was reanalyzed. **(c-d)** Histograms of the current conductance of TOM20-Flag-SVG28 **(c)** and TOM20-Flag-SVG28D2 **(d)**. Peak-fitting curves are plotted for peaks that can be fitted with a Gaussian function **(e-f)** I-V curves of TOM20-Flag-SVG28 **(e)** and TOM20-Flag-SVG28D2 **(f)**

To further characterize these pore structures, we analyzed their single-channel properties. TOM20-Flag-SVG28 showed an ionic conductance (*I/V*) of 0.3 nS, while TOM20-Flag-SVG28D2 showed 0.4 nS, with a subpopulation of 1.1 nS in 1 M KCl (**Fig. 2c-d**). Based on the relationship between pore size and pore conductance in natural β-barrel proteins ^30^, SVG28-Flag-TOM20 is estimated to form 5-mer pores composed of 10 β-strands with a pore diameter of 1.0 nm. Similarly, TOM20-Flag-SVG28D2 is estimated to form approximately 5-mer pores composed of 10.6 β-strands with a 1.1 nm pore diameter, while the 1.1 nS subpopulation corresponds to approximately 7-mer pores comprising 13.6 β-strands and a pore diameter of 1.7 nm, having sufficient pore sizes to transport protons.

Furthermore, current-voltage (*I-V)* curves exhibited an asymmetry with a higher current intensity at negative voltages (**Fig. 2e-f**). This is due to charge asymmetry in the SVG28 pore, wherein one terminus is positively charged and the other is negatively charged ^30^. This demonstrates that TOM20-Flag-SVG28 and TOM20-Flag-SVG28D2 retain the defining structural and biophysical characteristics of the native SVG28 pore.

### Localizing pore-forming synthetic peptides to mitochondria

To test whether the SVG28 peptides could manipulate organelle properties by forming a pore at a targeted location, DNA constructs encoding TOM20-Flag-SVG28, TOM20-Flag-SVG28D2, or TOM20-Flag as a control were generated (**Fig. 3a**). When each of these was transfected in HeLa cells, fluorescence imaging of these cells after immunostaining using a Flag antibody indicated all three constructs were successfully targeted to mitochondria (**Fig. 3b**). Strikingly, cells expressing Tom20-SVG28 or Tom20-SVG28D2, but not a control Tom20-Flag, exhibited altered mitochondrial morphology (**Fig. 3b**); while mitochondria in this cell type generally take tubular shape, SVG28- or SVG28D2-expressing cells showed more spherical mitochondria. (**Fig. 3b-c**). This result suggests that the pore-forming peptides SVG28 and SVG28D2 are capable of altering mitochondrial morphology in living cells when expressed on the mitochondrial outer membranes.

**Figure 3.**
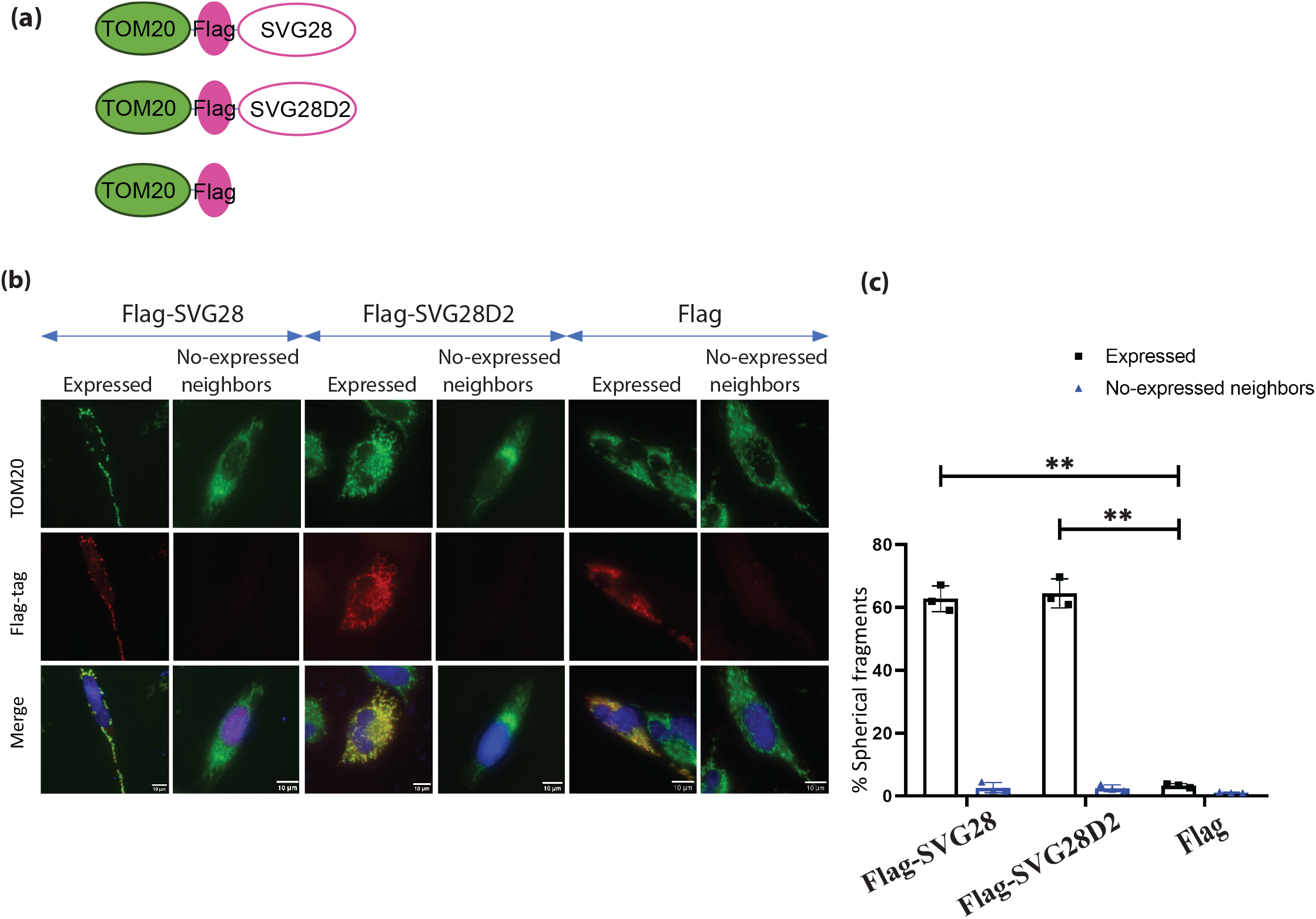
SVG28 and SVG28D2 located in mitochondrial membrane. **(a)** Illustration of the domains of the proteins **(b)** Representative images of mitochondria in HeLa cells labelled with Flag, Flag-SVG28, and Flag-SVG28D2. Scale bars, 10 μm **(c)** Graphs of the percentage of mitochondrial fragmentation. The means ± SD are shown (n=3). The P-values were obtained using a two-tailed paired Student’s t-test. Significance values are **p<0.01

### Functional remodeling of mitochondria

To evaluate the impact of mitochondrial targeting of SVG28 and SVG28D2 on mitochondrial properties, we constructed TOM20-Flag-SVG28, TOM20-Flag-SVG28D2, or TOM20-Flag, each of which was fused to a Halo tag via internal ribosomal entry site (IRES) (**Fig. 4a**). After expressing these constructs, we measured two major functions of mitochondria: membrane potential and ATP production using their fluorescence indicators (**Fig. 4b**).

**Figure 4.**
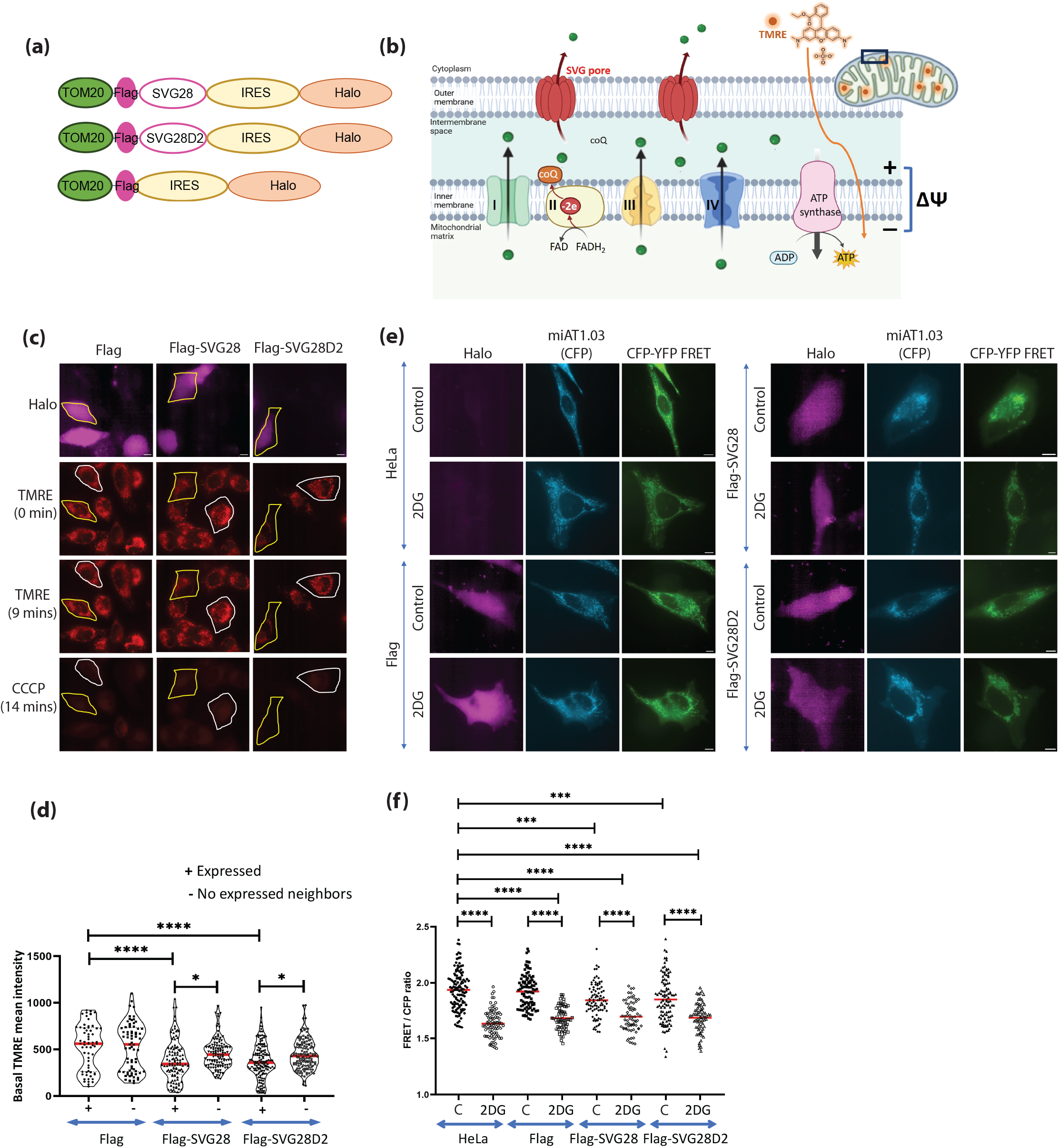
Effect of pore membrane on functional aspects of mitochondria. **(a)** Illustration of the domains of the proteins **(b)** Illustration of mitochondrial potential and ATP synthesis **(c)** Fluorescence images of TMRE in HeLa cells expressing Flag, Flag-SVG28, and Flag-SVG28D2 at 0 min, 9 mins before CCCP treatment, and after CCCP treatment of 14 mins. Scale bars, 10 μm **(d)** Graphs of TMRE mean intensity at mitochondria. The P-values were obtained using one-way ANOVA with Tukey’s post hoc analysis. Significance values are *p<0.05 and ****p<0.0001 **(e)** Representative FRET images of miAT1.03 along with TOM20-Flag, TOM20-Flag-SVG28, and TOM20-Flag-SVG28D2. Scale bars, 10 μm **(f)** Graphs of FRET/CFP ratio value. The P-values were obtained using one-way ANOVA with Tukey’s post hoc analysis. Significance values are ***p<0.001 and ****p<0.0001

We first assessed mitochondrial membrane potential in HeLa cells expressing either SVG28, SVG28D2, or Flag control. To measure membrane potential, we employed tetramethyl rhodamine methyl ester (TMRE). Cells were incubated with TMRE for 30 minutes before live-cell imaging. TMRE fluorescence intensity was lower in cells expressing SVG28 and SVG28D2 than in those expressing a control construct. carbonyl cyanide m-chlorophenyl hydrazone (CCCP) treatment resulted in complete loss of TMRE signal in all conditions (**Fig. 4c**). To evaluate the membrane potential, we quantified TMRE fluorescence intensity in mitochondria of expressing versus non-expressing cells. The result showed that the mitochondrial membrane potential was decreased by roughly 30% with the expression of pore-forming membrane peptides SVG28 and SVG28D2 (**Fig. 4d**).

We next measured mitochondrial ATP levels using mitAT1.03, an organelle-targeted Forster resonance energy transfer (FRET)-based biosensor of ATP ^33^. Methods for measurement and analysis of CFP-to-YFP FRET were described previously ^34-37^. FRET efficiency was determined by calculating the FRET signal normalized to CFP intensity. Expression of SVG28 and SVG28D2 resulted in a decrease in the FRET signal by roughly 10%, compared to the expression of the control construct (**Fig. 4e-f**). As a control, 2-deoxy-D-glucose was used to inhibit glycolysis and reduce ATP production. The 2DG treatment caused a strong decrease in the FRET signal in all three cases. This finding suggests that expression of pore-forming peptides leads to a reduction in ATP synthesis, likely due to the waning membrane potential.

### Mitochondrial inner membrane integrity

Given the observed reductions in mitochondrial membrane potential and ATP synthesis, we hypothesized that these effects might result from the disintegration of inner membranes due to compromise to their outer membrane. If this is the case, then this should lead to the release of matrix-resident proteins (**Fig. 5a**). To test this possibility, we co-expressed mitochondria matrix protein Subunit 9 (Su9) of mitochondrial ATPase, that was fused to mCherry with either TOM20-Flag-SVG28, TOM20-Flag-SVG28D2, or TOM20-Flag. Fluorescence imaging of Su9-mCherry revealed no significant difference in the intensity or subcellular distribution between the peptide-expressing and control cells (**Fig. 5b-c**). These results indicate that the peptides did not cause any detectable leakage of soluble matrix proteins into the cytosol. Combined with the characterization *in vitro*, it is highly likely that SVG28 peptides formed a pore on the outer membranes, which permeated protons in the intermembrane space of mitochondria, thereby dissipating the proton gradient across the inner membranes.

**Figure 5.**
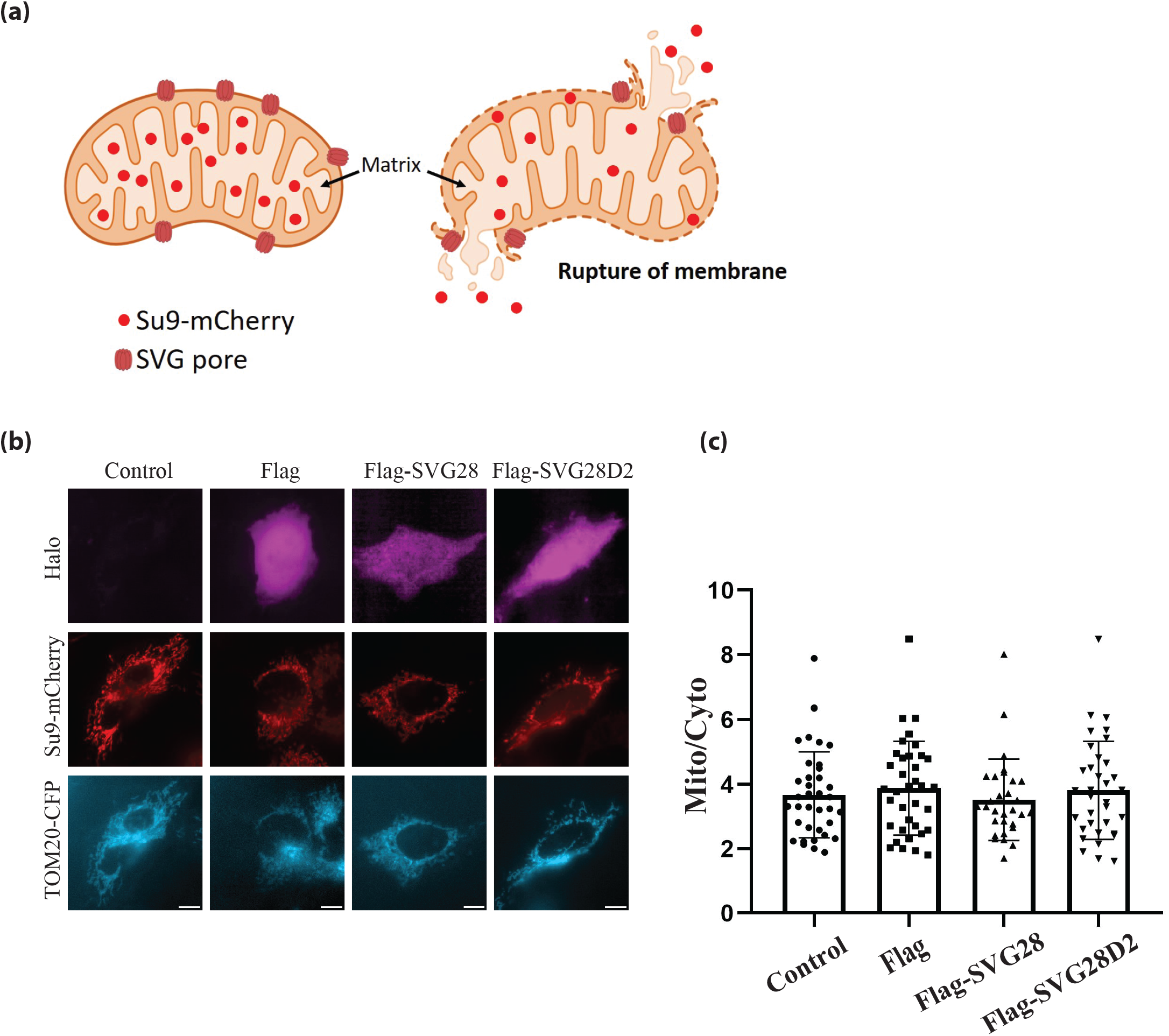
Mitochondrial matrix protein did not release to cytosol. **(a)** Illustration of the presence of SVG28 or SVG28D2 (SVG pore) at the outer mitochondrial membrane, with Su9-mCherry which localized to the mitochondrial matrix, under condition of intact mitochondria or rupture of mitochondrial membrane **(b)** Representative images of Su9-mCherry showing Su9-mCherry localization to matrix of mitochondria co-expressed with TOM20-Flag, TOM20-Flag-SVG28, or TOM20-Flag-SVG28D2. Scale bars, 10 μm **(c)** Plots illustrating the relative mitochondrial-to-cytosolic fluorescence intensity ratio. No statistically significant differences were observed (p> 0.05).

## Discussion

In this study, we demonstrate a unique strategy for precise manipulation of mitochondrial morphology and function by targeting engineered pore-forming peptides, SVG28 and SVG28D2, to the mitochondrial outer membrane via fusion with TOM20. Our results show that these peptides localize to mitochondria in HeLa cells, inducing mitochondrial fragmentation accompanied by decreased membrane potential and ATP production, while maintaining matrix compartmentalization. This highlights selective modulation of mitochondrial membrane integrity without catastrophic membrane rupture.

Interestingly, the fusion of SVG28 to the transmembrane domain of TOM20 appears to enhance the stability of SVG28. While TOM20 is conventionally used as a targeting sequence for the outer membrane localization of exogenous proteins ^38-40^, its demonstrated structural compatibility with SVG28 may also characterize TOM20 as a stabilizing scaffold that reduces proteolytic degradation and/or maintains membrane insertion competence.

Channelrhodopsins are light-gated cation channels composed of single-chain, seven-transmembrane helix proteins, typically 40–77 kDa in size, and have been targeted to mitochondria to achieve temporal control of ion flux and membrane potential ^41-43^. However, their relatively large and complex architecture may limit broader applicability within the mitochondria and also in other organelles. These challenges underscore the need for smaller, more stable, mitochondria-adapted tools to reliably modulate mitochondrial physiology without the drawbacks associated with native channelrhodopsin constructs ^41; 42^ By contrast, SVG28 pores are composed of ∼3 kDa monomers, each forming two transmembrane domains, whose minimal architecture may facilitate more efficient integration into organelle membranes.

In the future, strategies could be developed to regulate the gating of SVG28 pores, such as engineering peptide variants responsive to chemical inducers, small molecules, or light. For example, light-controllable SVG28 pores could be achieved by incorporating photo-switchable amino acids or optogenetic modules. In conclusion, we present a synthetic biology approach to directly manipulate mitochondrial integrity and function by leveraging 28–amino acid peptides rationally designed de novo to self-assemble into nanoscale pores within biological membranes. This platform provides a versatile foundation for advancing applications in cell biology and nanotechnology. Because these synthetic molecular tools are genetically encoded, they can be introduced readily into target cells, tissues, and organisms. Together, these unique features open opportunities for organelle-targeted therapies and the creation of synthetically engineered organelles.

## Methods

### Plasmid construction

Plasmids of TOM20-Flag, TOM20-Flag-SVG28, and TOM20-Flag-SVG28D2 were cloned into the C1 vector using Gibson assembly. The gene encoding IRES-Halo was subcloned into the C1 vector of TOM20-Flag, TOM20-Flag-SVG28, and TOM20-Flag-SVG28D2. All plasmids were cloned or subcloned using Gibson assembly, transformed into DH5α, and confirmed by DNA sequencing.

### Cell culture

HeLa cells (ATCC CCL-2) were maintained in Dulbecco’s Modified Eagle Medium (DMEM; Corning, 10-013-CV) supplemented with 10% fetal bovine serum (FBS; Sigma-Aldrich, F6178), and 1% penicillin-streptomycin (Pen-Strep; Thermo Fisher Scientific, 15140163). Cells were cultured at 37°C in a humidified incubator under 5% CO□.

### MD simulations

The 7-mer structure of TOM20-Flag-SVG28D2 was modeled using the AlphaFold3 ^31^ server with 40 oleic acid molecules to mimic membranous environment. Due to the undesirable predicted topology for TOM20-Flag-SVG28, its 7-mer structure was modeled using homology modeling ^44^ with TOM20-Flag-SVG28D2. Each structure was embedded in DOPC:Chol. (2:1 mol:mol) membranes with 1 M KCl using the CHARMM-GUI membrane builder ^45^. The systems were initially equilibrated using the protocol provided by the CHARMM-GUI, followed by an additional 500 ps of equilibration, and subsequently simulated for 200 ns with the CHARMM36m force field. All simulations were executed with GROMACS-2024.1 ^46^.

### Cell-free synthesis

DNA templates for CFPS were synthetic genes provided by Integrated DNA Technologies, USA. Purefrex2.0 (GeneFrontier, Japan) ^47^ was used for CFPS. Synthesis was performed at 37LJ for 6 hours. A reaction mixture containing 1 ng/µL DNA, 20 µL of solution I, 2 µL of solution II, 4 µL of solution III, and 10 µL of nuclease-free water was prepared. After the translation, the peptides were desalted and purified with GL-Tip SDB (GL Sciences, Japan). Matrix-assisted laser desorption/ionization time-of-flight (MALDI-TOF) mass spectrometry was performed using the Autoflex Speed MALDI-TOF instrument (Bruker Daltonics Inc., Billerica, MA, USA) in linear positive mode. PEG6000 was used for external mass calibration. Lyophilized peptides obtained from 10 μL of the reaction mixture were dissolved in 3 μL of TA30 solution (acetonitrile:0.1% trifluoroacetic acid = 3:7, v/v). A matrix solution was prepared by dissolving 100 μg of α-cyano-4-hydroxycinnamic acid (CHCA) in 100 μL of TA30. The peptide sample and matrix were mixed at a 1:2 (v/v) ratio, and 1 μL of the mixture was spotted onto the MALDI target plate.

### Liposome-added cell-free synthesis

Liposomes were prepared using the gentle hydration method. First, DOPC and DOPG dissolved in chloroform at 1-to-1 molar ratio were placed in a glass microtube and evaporated. The lipid films were hydrated with 500 µL of 0.5 M glucose for 3 hours at 40LJ. After the hydration, 200 µL of 0.5 M glucose and 100 mM HEPES (pH 7.4) were added to 200 µL of liposome suspension. The liposome solution was extruded using an extruder (Avanti Polar Lipids, USA) equipped with a 1.0 µm-pored membrane (Whatman, UK). A 20 µL of the CFPS reaction mixture described above was prepared with the addition of 10 µL of 10 mg/ml liposome solution, for a total volume of 30 µL. After the CFPS, the reaction mixture was left at 4LJ for approximately one week. Then, 150 µL of 50 mM HEPES was added to the PURE reaction solution, which was subsequently centrifuged at 20,000 xg for 5 min. The supernatant was carefully discarded after centrifugation. Another 150 µL of 50 mM HEPES was added to the solution, and then it was centrifuged again. The supernatant was completely removed, and the pellet was resuspended in 10 µL of 50 mM HEPES. A mixture of 10 µL of the liposome solution and 190 µL of 500 mM glucose and 50 mM HEPES was extruded using an extruder equipped with a 0.1 µm-pored membrane (Whatman, UK).

### Channel current measurement

As shown in our previous reports, we prepared planar bilayer lipid membranes (BLMs) using the droplet contact method within a microdevice, with two 4.0 mm-diameter wells and a 100-µm-diameter pore for the droplet contact site in a 5 µm-thick parylene C (polychloro-*p*-xylylene, Parylene Japan, Japan) film (**Supplementary Fig. 4**). Each well had a through-hole at the bottom containing Ag/AgCl electrodes. Two lipid monolayers containing the aqueous droplets came into contact through the small hole in the parylene film, forming lipid bilayers.

First, 2.5 µL of 10 mg/ml DOPC:Cholesterol= 2:1 (mol:mol) in *n*-decane was added to both chambers. Then, 23 µL of solution containing 1 mg/ml liposome, 250 mM glucose, 25 mM HEPES, 1 M KCl, and 10 mM MOPS was added to the voltage-applied chamber, and 23 µL of solution containing 1 M KCl and 10 mM MOPS was added to the grounded chamber. The channel current was monitored using a JET patch-clamp amplifier (Tecella, USA). Signals were detected with a 4 kHz low-pass filter at a sampling frequency of 20 kHz. A constant voltage of +100 mV was applied to the recording chamber, and the ground side chamber was grounded. Data were analyzed using Clampfit 11.3 (Molecular Devices, USA), Excel (Microsoft, USA), and OriginPro 2022 (OriginLab Corporation, USA).

To quantify pore-forming ability, current signals were classified into stable or unstable pore-formation signals ^30; 48^. Stable pore formation is characterized by a sharp increase in signal rise in less than 10 ms and a 95% confidence interval of the pore current value that is 10 times less than the baseline, or by a sharp increase in signal rise in less than 10 ms with staircase-like current-time traces and a 95% confidence interval of the pore current value that is 10 times less than the baseline for each step (**Supplementary Fig. 5a**). Unstable pore formation is characterized by a sharp increase in signal rise in less than 10 ms and a 95% confidence interval of the pore current value that is 10 times greater than the baseline, or by a blunt increase in signal rise in more than 10 ms (**Supplementary Fig. 5b**). Signals longer than 5 s and greater than 0.2 nS were used for analysis. Signal distributions were calculated using n = 20 data for one measurement device.

### Mitochondrial Localization Assay

HeLa cells per well were seeded per well into poly-D-lysin-coated 8-well glass chamber slides (Sigma, P6407-5MG, Cellvis, C8-1-N). Cells were transfected the following day with plasmids of TOM20-Flag, TOM20-Flag-SVG28, and TOM20-Flag-SVG28D2 using FUGENE HD (Promega, E2311), according to the manufacturer’s protocol. After 16 hours of transfection, cells were fixed and immunostained using primary antibodies against Flag (mouse, Millipore Sigma, F1804) and TOM20 (rabbit, Cell Signaling, 42406S). For imaging, an Eclipse Ti inverted fluorescence microscope (Nikon) equipped with a x60 oil-immersion objective lens was used.

### Mitochondrial membrane potential measurement

To assess mitochondrial membrane potential, HeLa cells were seeded and transfected as described above. After 16 hours, cells were incubated with tetramethyl rhodamine ethyl ester (TMRE; Thermo Fisher) at a final concentration of 7 nM for 30 minutes at 37°C before live imaging. TMRE fluorescence was used as a measurement of mitochondrial inner membrane potential. As a positive control for membrane potential dissipation, cells were treated with the proton ionophore carbonyl cyanide m-chlorophenyl hydrazone (CCCP) at a final concentration of 10 μM. Live-cell imaging was performed for 10 minutes following TMRE staining, followed by an additional 15 minutes of imaging during CCCP treatment using an Eclipse Ti inverted fluorescence microscope equipped with a x60 oil-immersion objective.

### Mitochondrial ATP synthesis measurement

To evaluate ATP production in mitochondria, a mitochondria-targeted FRET-based ATP biosensor, mitAT1.03 was used with CFP excitation light and collecting emission with YFP emission filter at 37°C. HeLa cells were seeded in 8-well glass-bottom chambers coated with poly-D-lysin and then were co-transfected with mitAT1.03 and either TOM20-Flag-IRES-Halo, TOM20-Flag-SVG28-IRES-Halo, or TOM20-Flag-SVG28D2-IRES-Halo using FuGene-HD. After transfection, live-cell imaging was performed at 37°C using excitation at the CFP wavelength and capturing emission through a YFP filter to measure FRET signal. For a control condition, cells were treated with 25 mM 2-deoxy-D-glucose (2-DG) to inhibit glycolysis and reduce ATP production. Fluorescence intensities from the CFP and FRET channels were quantified using FIJI (ImageJ), and the FRET/CFP ratio was calculated for each condition. Data represent the average of at least three biologically independent experiments.

### Su-9 at matrix mitochondria release

To determine the function of pore membrane for protein release, Su9-mCherry was co-transfected with TOM20-Flag-IRES-Halo, TOM20-Flag-SVG28-IRES-Halo, or TOM20-Flag-SVG28D2-IRES-Halo in HeLa cells using FuGene-HD after cells seeded in 8-well glass-bottom chambers coated with poly-D-lysine. After transfection of 16 hours, live-cell imaging was performed at 37°C with 5% COL□. Fluorescence intensities of Su9-mCherry at mitochondria and cytosol were quantified using FIJI (ImageJ), and the Mitochondria/Cytosol ratio (Mito/Cyto ratio) was calculated for each condition. Data represent the average of at least three biologically independent experiments.

### Statistical analyses

All data are mean ± SD, as indicated in the legends. Data for each condition were obtained from at least three independent biological replicate experiments. The two-tailed paired Student’s t-test was used to measure differences between the two groups. For multiple-group comparisons, one-way ANOVA was applied under the assumption of normal distribution. Detailed statistical methods are described in the individual figure legends. These statistical analyses were performed by using GraphPad Prism 8. A value of *p*<0.05 was considered significant.

## Acknowledgement

We thank members of the Inoue and Kawano laboratories for technical assistance and valuable discussions. In addition, we are grateful for the initial support from the Global Innovation Research (GIR) Program organized by the Tokyo University of Agriculture and Technology.

## Author Contributions

TI and RK conceived the idea. NTHN initiated the project. NTHN, RK, and TI designed the experiments. NTHN performed the experiments and analyzed the data under the supervision of TI. SF and NT performed experiments and prepared the draft of the sections on MD simulations and electrophysiological measurements under the supervision of RK. NTHN wrote all the remaining manuscript in consultation with RK and TI. All the authors contributed to the final version of the manuscript.

## Data Availability

The datasets generated during this study are available from the corresponding author upon request. Source data are provided with this paper.

