## Supplementary material for "Self-Assembling Synthetic Peptides Remodel Mitochondrial Properties via Nanopore Formation": Supp figures 1-5

**Supplementary Figure 1. Quantitative analysis of MD simulation and synthesis confirmation of TOM20-Flag-SVG28**

**(a)** Conformational changes during 200 ns simulations of TOM20-Flag-SVG28 as measured by root mean square deviation (RMSD) from the initial model

**(b)** Root mean square fluctuation (RMSF) of each protein Cα atom of the last 50 ns simulations

**(c)** MALDI-TOF/MS mass spectra of TOM20-Flag-SVG28 expressed using cell-free synthesis

**Supplementary Figure 2. TOM20-Flag-SVG28D2 conjugate**

**(a)** TOM20-Flag-SVG28D2 structures predicted by *in silico* methods. TOM20 in the transmembrane region is shown in purple; TOM20 outside the membrane in magenta; flag tag in orange; the linkers in green; SVG28D2 in cyan

**(b)** TOM20-Flag-SVG28D2 pore structures after 200 ns simulation in DOPC:Chol membrane. DOPC and cholesterol molecules are shown in yellow

**(c)** MALDI-TOF/MS mass spectra of TOM20-Flag-SVG28D2 expressed using cell-free synthesis

**Supplementary Figure 3. Channel current of cell-freely synthesized TOM20-Flag-SVG28 and TOM20-Flag-SVG28D2 without liposomes**

**(a-b)** Observed membrane interaction signals of TOM20-Flag-SVG28 **(a)** and TOM20-Flag-SVG28D2 **(b)**. They were synthesized using cell-free synthesis without liposomes

**Supplementary Figure 4. Microdevice for channel current measurement**

**(a)** A microdevice has two sample chambers separated by a separator region and Ag/AgCl electrodes embedded in the bottom of the sample chamber

**(b)** A 3D image of a microdevice. Parylene film is sandwiched between two separators embedded in the square grooves

**(c)** A bilayer lipid membrane is formed in the hole of the parylene film

**(d)** In the droplet contact method, two lipid monolayers containing aqueous droplets come into contact to form a lipid bilayer.

Figures were adapted from S. Fujita *et al., ACS Nano* (2023), licensed under a CC BY 4.0.

**Supplementary Figure 5. Classification of current signals**

**(a-b)** Detailed criteria for classifying signals for stable **(a)** and unstable **(b)** pore formation. Signals were classified according to the time it takes to rise, the current width after rising, and the presence of transition states. Figures were adapted from S. Fujita *et al., ACS Nano* (2023), licensed under a CC BY 4.0.

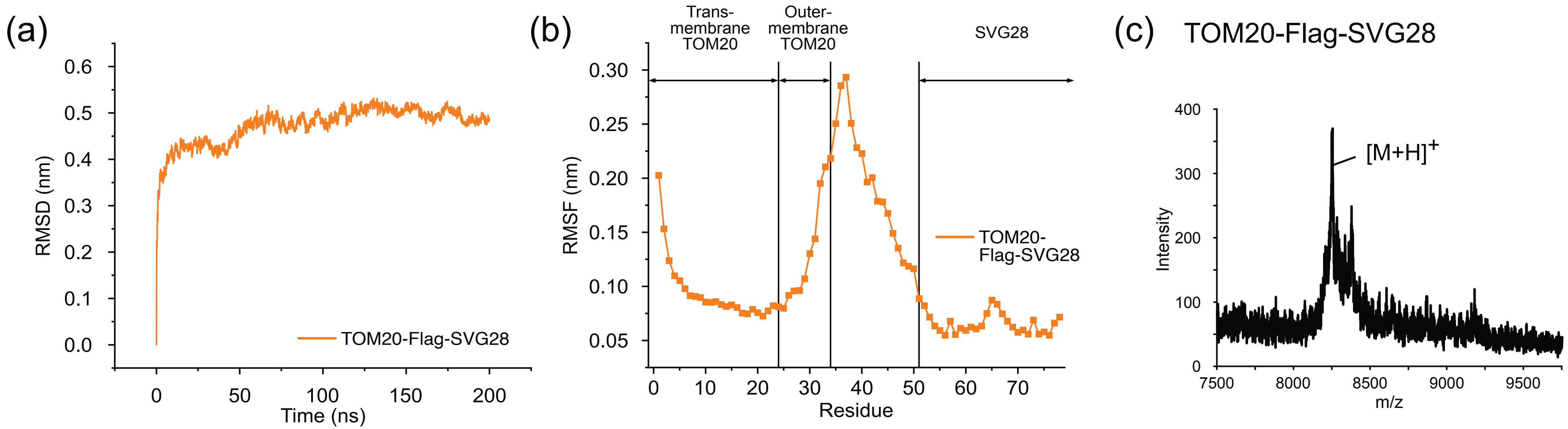


**Supplementary Figure 1. Quantitative analysis of MD simulation and synthesis confirmation of TOM20-Flag-SVG28.**

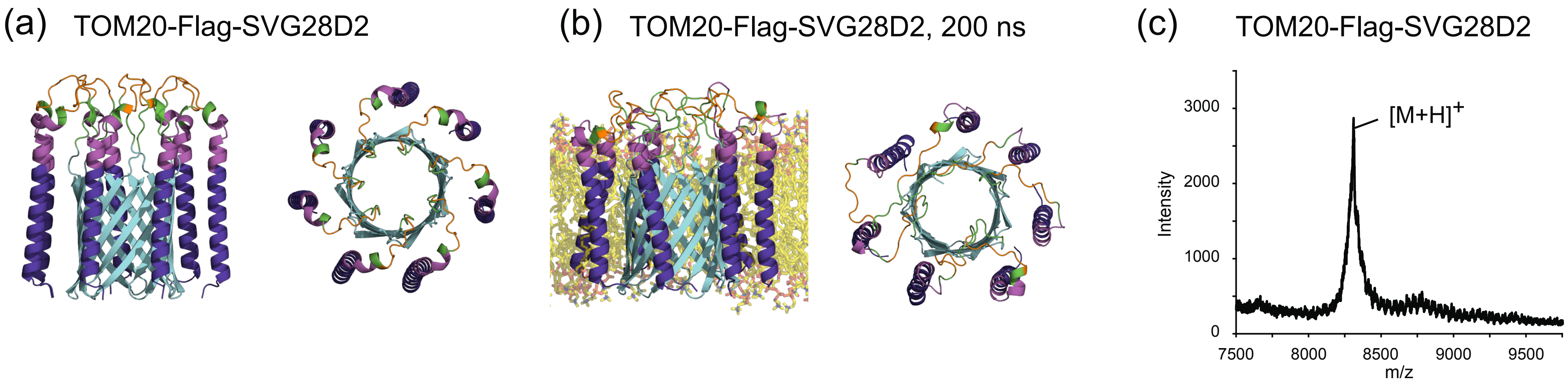


**Supplementary Figure 2. TOM20-Flag-SVG28D2 conjugate.**

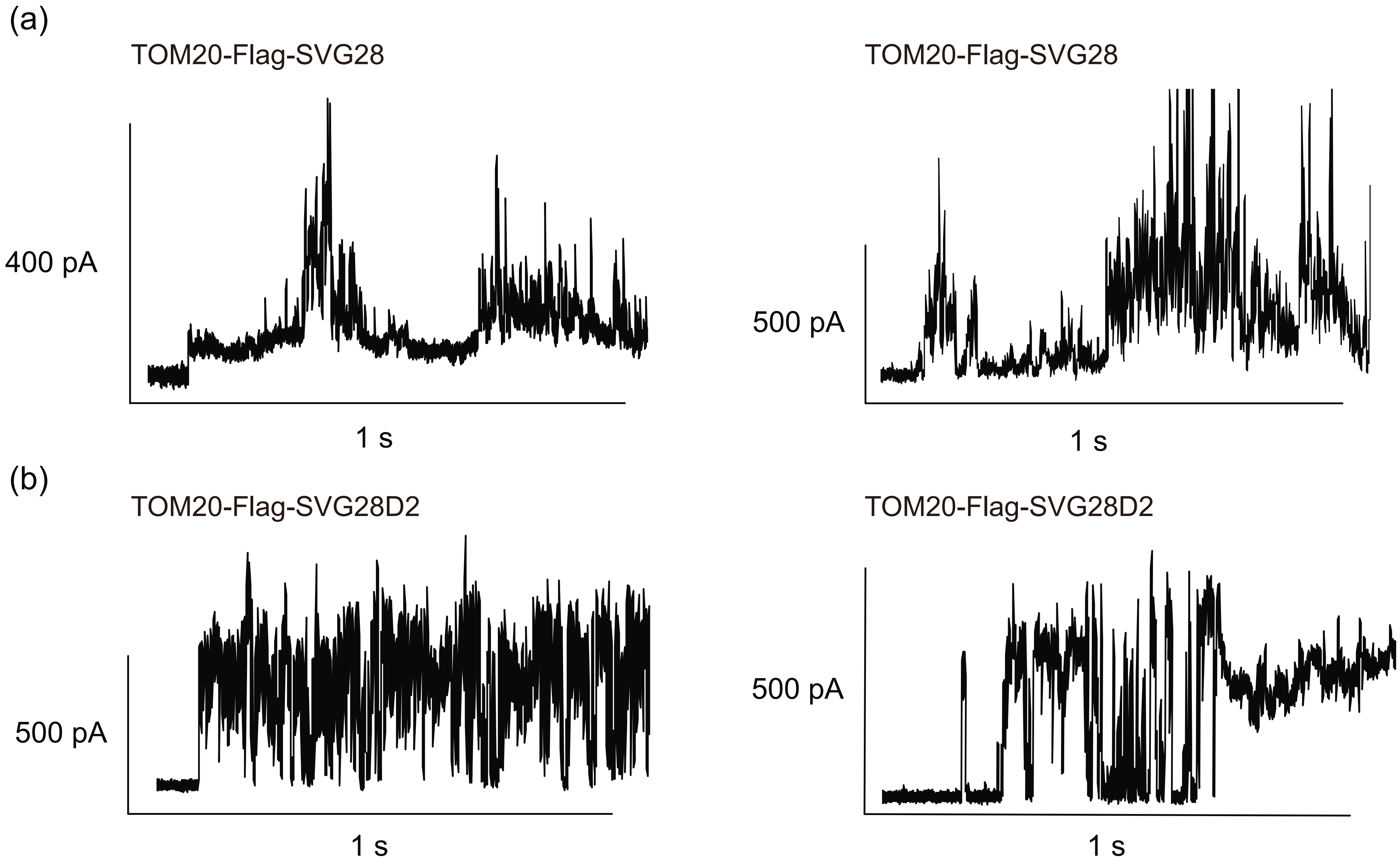


**Supplementary Figure 3. Channel current of cell-freely synthesized TOM20-Flag-SVG28 and TOM20-Flag-SVG28D2 without liposomes**

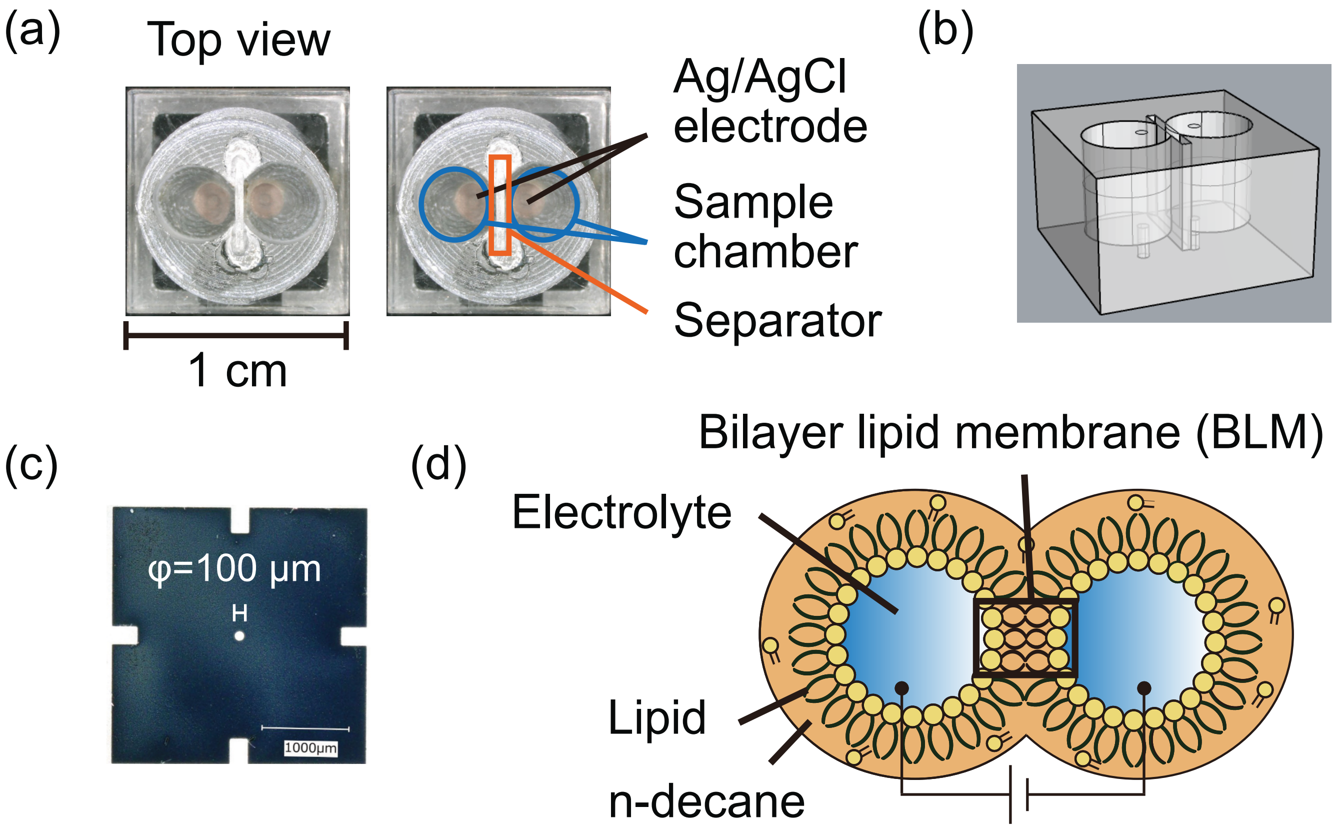


**Supplementary Figure 4. Microdevice for channel current measurement**

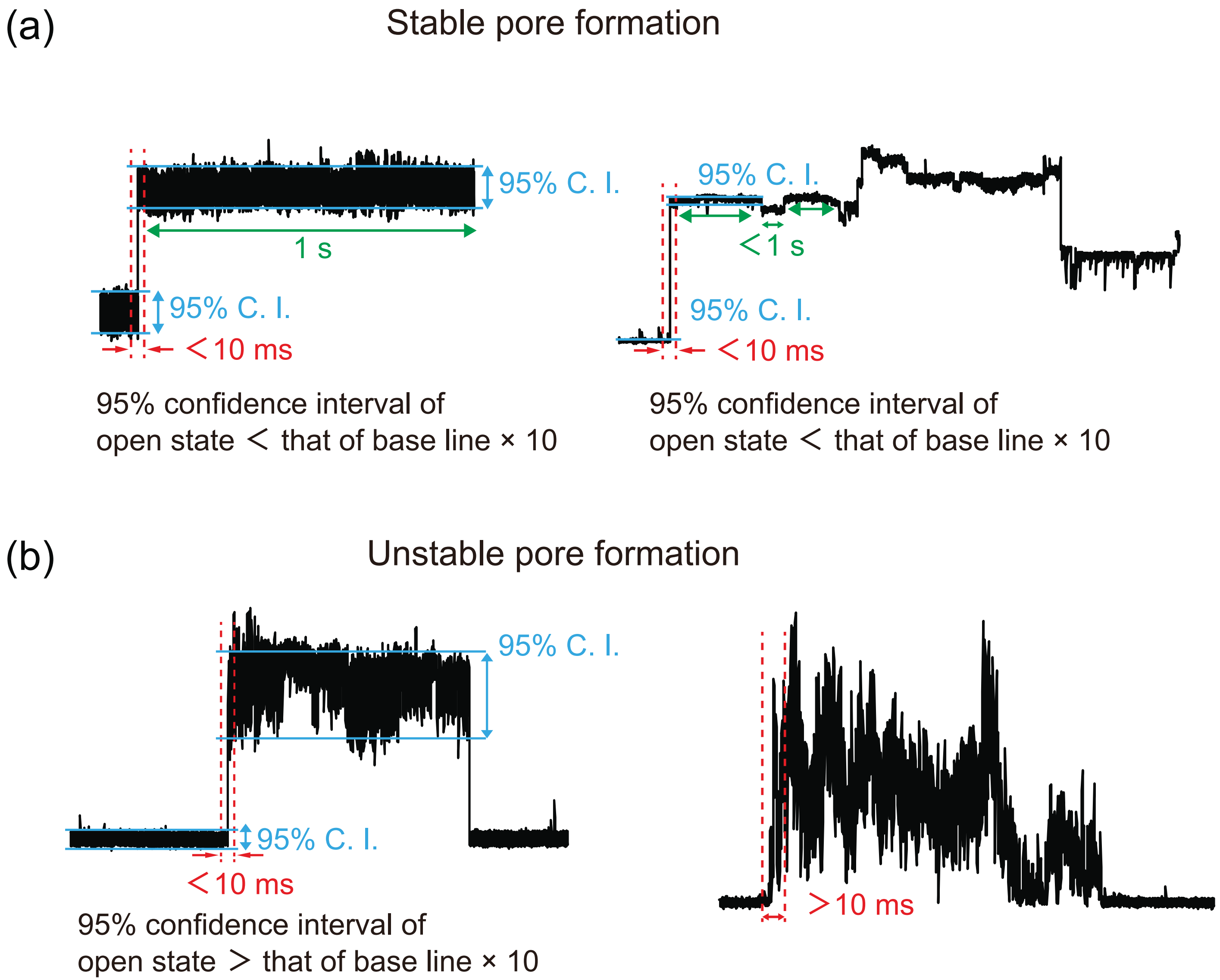


**Supplementary Figure 5. Classification of current signals**
